# Improving the temporal accuracy of functional Magnetic Resonance Imaging

**DOI:** 10.1101/073437

**Authors:** Niels Janssen, Juan A. Hernández-Cabrera, Laura Ezama Foronda

## Abstract

A major drawback of functional Magnetic Resonance Imaging (fMRI) concerns the lack of temporal accuracy of the measured signal. Although this limitation stems in part from the neuro-vascular nature of the fMRI signal, it also reflects particular methodological decisions in the fMRI data analysis pathway. Here we show that the temporal accuracy of fMRI is affected by the specific way in which whole-brain volumes are created from individually acquired brain slices. Specifically, we show how the current volume creation method leads to whole-brain volumes that contain within-volume temporal distortions and that are available at a low temporal resolution. To address these limitations, we propose a new framework for fMRI data analysis. The new framework creates whole-brain volumes from individual brain slices that are all acquired at the same point in time relative to a presented stimulus. These whole-brain volumes contain no temporal distortions, and are available at a high temporal resolution. Statistical signal extraction occurs on the basis of a novel time point-by-time point approach. We evaluated the temporal characteristics of the extracted signal in the standard and new framework with simulated and real-world fMRI data. The new slice-based data-analytic framework yields greatly improved temporal accuracy of fMRI signals.

---

The changes in blood oxygen concentration that result from neural activity are detected by functional Magnetic Resonance Imaging (fMRI) in the form of changes in the Blood Oxygen Level Dependent (BOLD) signal (Ogawa et al., 1990). The BOLD signal is observed in T2* weighted imaging runs and represents changes in the magnetic field strength as a function of time. It has a characteristic shape that involves an initial dip, a dispersed peak, and a post-stimulus undershoot (Hu et al., 1997; Menon et al., 1998). The BOLD signal is sluggish and peaks about 5 seconds after the onset of neural activity. The precise shape of the BOLD signal is of crucial importance in event-related fMRI studies that examine BOLD signal dynamics in response to the presentation of stimuli (Friston et al., 1998; Josephs et al., 1997), in studies of functional connectivity, where the connection strength between different areas of the brain is established on the basis of the overall similarity in their BOLD signal dynamics (Biswal et al., 1995; Rissman et al., 2004), and in studies of neuro-vascular coupling, where BOLD signal dynamics are used to make claims about the metabolic implications of neuronal activity (Attwell & Iadecola, 2002; Hillman, 2014; Logothetis & Wandell, 2004; Uludağ & Uğurbil, 2015). A crucial question for current fMRI studies is therefore related to the temporal accuracy and resolution by which the fMRI BOLD signal can be estimated from the data. We will first discuss the details of fMRI data acquisition and outline why the current method for creating whole-brain volumes produces a lack of temporal accuracy and poor temporal resolution. We will then discuss our new proposal for improving the temporal aspects of fMRI BOLD signal extraction.

## fMRI data acquisition and volume creation

One of the most common forms of data acquisition in fMRI experiments is one where the data are obtained by progressively taking small samples from different parts of the brain. Specifically, when an animal or human subject is positioned in the MRI scanner, fMRI data is acquired in the form of two-dimensional planes of data points called *slices*. Each slice covers a small, specific and unique portion of the brain. To provide whole-brain coverage, typically more than 30 slices are required. Different from other techniques such as EEG and MEG (e.g., Nunez & Srinivasan, 2006), current MRI technology does not permit the instantaneous sampling of all data points on all slices in the entire brain. Instead, the most common form of data acquisition involves the progressive sampling of single slices in sequence (Cohen & Weisskoff, 1991) or of a few slices in parallel (Moeller et al., 2010). Slices are acquired at well-defined points in time. Currently, the time between successive samples is on the order of several tens of milliseconds. This means that with typical MRI parameters (i.e., the TR), the time from sampling the first slice to the last slice in a whole-brain covering series takes around 3 seconds.

Formally, fMRI data *D* can be represented as a set of *m* slices *S* that are repeatedly sampled *n* times:

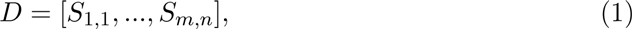

where each *S* is itself a two dimensional matrix of acquired fMRI signal intensities (not shown here). This data matrix of slices *D* is accompanied by a similar size *m × n* matrix of slice acquisition times *DT*.

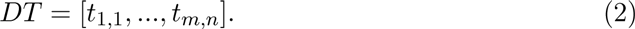

Under the assumption of a standard sequential slice acquisition scheme, each specific time point *t* (*a, b*) in this matrix can be determined by the following function:

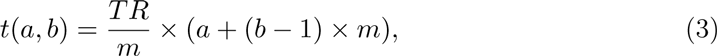

where *a* and *b* index the specific slice and acquisition number. Adjustments to this formula are required for more complex (i.e., interleaved) acquisition schemes. fMRI data can thus be represented as a long series of slices sampled at well-defined time points.

This method of progressively sampling a small area of the brain raises a fundamental problem for fMRI data analysis. Specifically, the problem is how to perform whole-brain analyses of the fMRI signal when at each point in time only data from a small part of the brain are available. The solution to this problem is not straightforward. At the moment, the only available solution is to create whole-brain volumes by temporally displacing the individually acquired slices (e.g., Friston et al., 1994). Specifically, according to this solution, sets of sequentially acquired slices that cover an entire brain are simply assumed to be acquired at the same moment in time. This moment in time is defined with respect to an arbitrarily chosen referent slice, for example the first or the middle slice in the set. This then creates series of whole-brain volumes, each of which contains a set of slices that are all assumed to be acquired at the moment in time when the referent slice was acquired (see Figure 1 for a graphical explanation).

**Figure 1:**
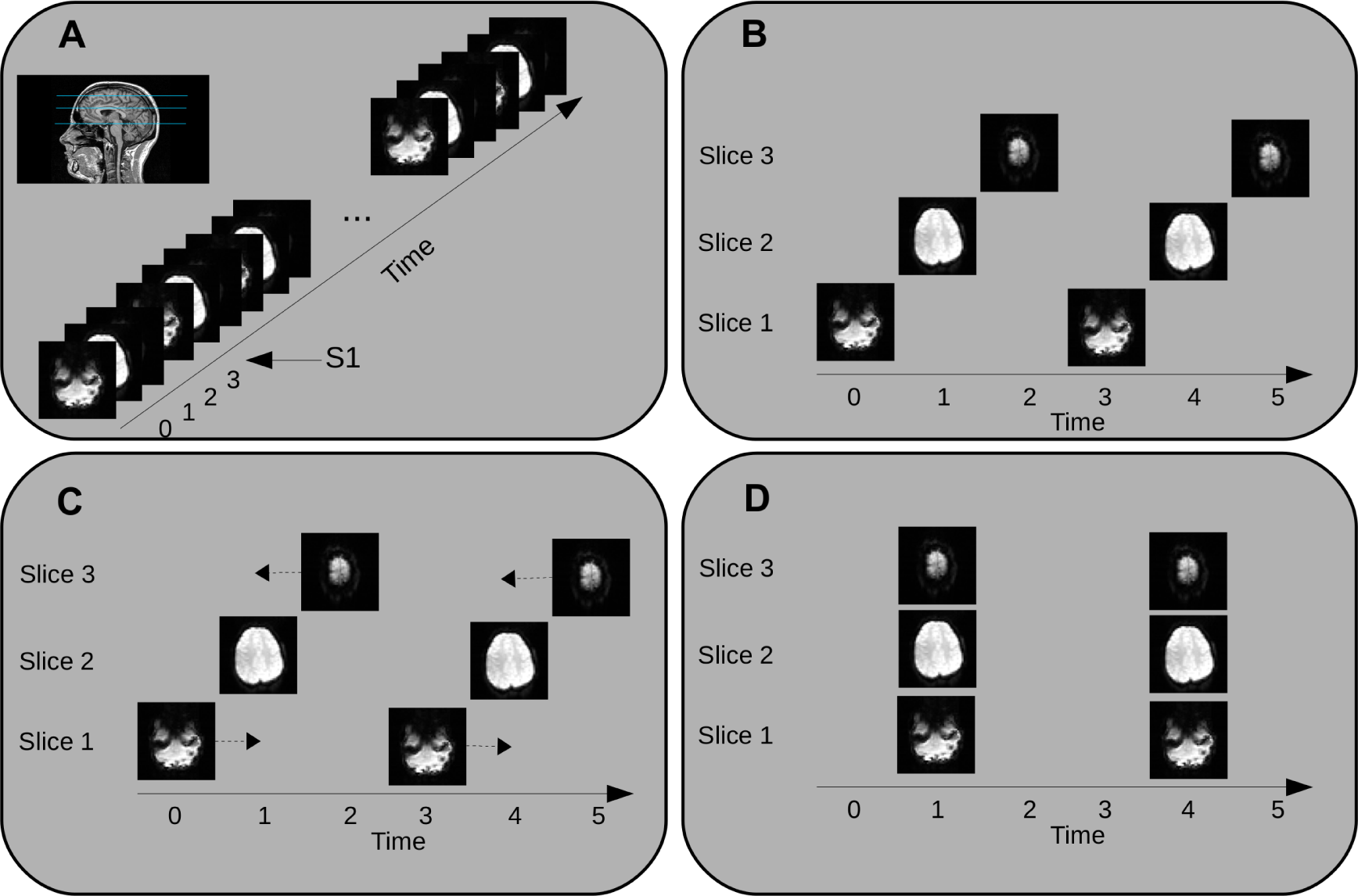
Current standard method for creating whole-brain volumes from raw fMRI data. Panel A shows an imaging run where a set of three slices are sequentially sampled at well defined points in time. Panel B reveals the same data, reorganized to illustrate that at no sampled time point information from the whole-brain is available, requiring data transformation. Panel C shows the standard solution, where slices are time-shifted to new positions in time (arrows indicate shift direction), using the middle slice as an arbitrary referent. Panel D shows the final transformed data, where whole-brain volumes are available every TR. Note how the final volumes contain slices acquired at different points in time, and how time points where data was sampled are no longer used.

Formally, the raw fMRI data *D* is transformed from *m × n* individual slices, each acquired at distinct point in time to a *n* size vector *D*′ of whole-brain volumes *V*_1_,…, *V_n_*:

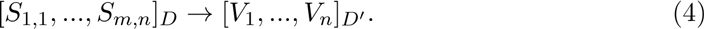

In this new formulation of the data *D*′, it is simply assumed that all slices within a given volume are acquired at the same point in time. Specifically, the vector of volume acquisition times is give by:

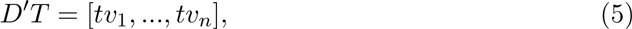

where each volume acquisition time *tv*(*v*) is determined by the function:

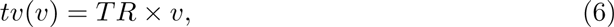

where *v* ranges from 1 to *n*. In a typical fMRI experiment, the *tv* values are on the order of seconds.

This method of creating whole-brain volumes by *time-shifting* individual slices leads to two major problems with respect to the temporal aspects of the fMRI signal. First, the method produces within-volume temporal distortions. Specifically, within this method, volumes are composed out of slices that were *not* acquired at the same moment in time. In the real-world terms, this means that voxels that are only a few mm apart in the brain could have been sampled seconds apart in time. This temporal distortion of the signal that is present within each volume causes the whole-brain volumes created within this method to be temporally imprecise. Note that this aspect of fMRI data acquisition is well-known, and is the reason for the existence of a data-analytic procedure called Slice Time Correction (STC; Henson et al., 1999; Sladky et al., 2011). This procedure attempts to account for the within-volume temporal distortion by making the data of slices within a volume more like those of the referent slice (e.g., by time shifting signals). However, although this may improve basic signal detection methods (Sladky et al., 2011), its impact on improving the temporal aspects of signal extraction remain uncertain. We further addressed this issue below.

The second problem with the aforementioned volume creation method concerns the temporal resolution by which the volumes are created. Specifically, the sampling frequency in the original data format *D* is often is on the order of several tens of milliseconds. Specifically, the sampling frequency of the original raw data is proportional to the ratio between the TR and the number of slices *m* (see Equation 3). By contrast, the sampling frequency of the transformed data *D*′ is generally on the order of seconds. Specifically, in this case the sampling frequency is directly proportional to the *TR* (see Equation 6). Note that this temporal resolution of the transformed data *D*′ is therefore one or two orders of magnitude below the actual temporal resolution by which data was sampled from the brain in *D*. Thus, even though data was sampled at a high temporal resolution, this temporal resolution is unfortunately lost in the transformed data *D*′.

To summarize, the current standard method creates whole-brain volumes from raw fMRI data by time-shifting individually acquired slices into whole-brain volumes. As we explained above, this leads to volumes that contain signals that are temporally inaccurate and that are of a low temporal resolution. These limitations on the temporal aspects of fMRI data are relevant for all current fMRI studies that rely on inferences based on the dynamics of the fMRI BOLD signal. In this light, we have created an alternative framework for fMRI BOLD signal extraction.

### Slice-based fMRI: An alternative method for volume creation

The key aspect of the new volume-creation method is that whole-brain volumes contain slices that are all acquired at the same point in time *relative to a presented stimulus*. The method places important constraints on the timing of stimulus presentation in an imaging run. In particular, stimuli have to be presented in-phase with the acquisition of the different slices that provide the whole-brain coverage. Volume creation in this framework relies on the combination of slices that were acquired during different stimulus presentations, but that were acquired at the same point in time relative to a stimulus. This method leads to the creation of whole-brain volumes that contain no temporal distortions and that have a temporal resolution equal to the sampling frequency (see Figure 2 for a graphical explanation). Given the importance of slices in this method, we will refer to this method of volume creation as *Slice-Based fMRI*.

**Figure 2:**
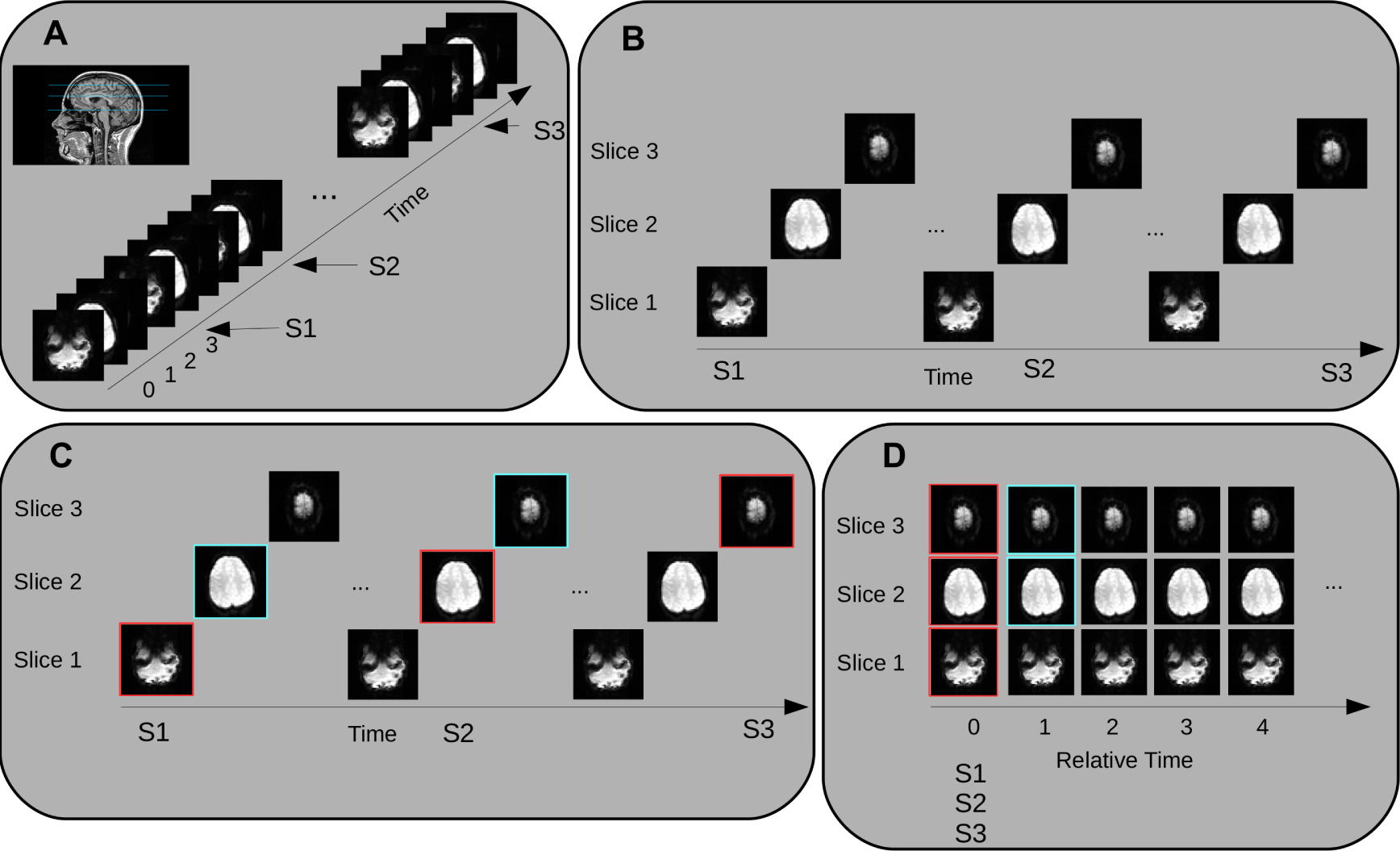
Slice-based method for creating whole-brain volumes from raw fMRI data. Panel A shows an imaging run where again three slices are sampled sequentially. Three stimuli S1, S2, and S3 of the same experimental class are presented during the run. Panel B shows that these stimuli are presented in-phase with slice acquisitions: S1 is presented in-phase with acquisition of slice 1, S2 with slice 2, and S3 with slice 3. Panel C shows how whole-brain volumes are created. Slices acquired at the same point in time relative to the onset of a stimulus can be combined (e.g., those highlighted in red and magenta). Panel D shows the final transformed data, where whole-brain volumes are available that only contain slices that are acquired at the same moment in time relative to a presented stimulus, and where whole-brain volumes are available at the sampling frequency (here TR/3).

Consider a set *P* of *m* stimuli [*p*_1_,…, *p_m_*], whose corresponding stimulus presentation times *P T* coincide precisely with the slice acquisition times determined by equation 3:

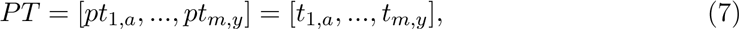

where *a,…, y* is chosen such that there is a sufficiently long time in between stimulus presentations. Note also that the number of stimuli is equal to the number of slices *m*. Next, we create *m* epochs *E*_1_,…, *E_m_* corresponding to each stimulus presentation. Each epoch has length Δ*t*. For a given epoch *E_j_* corresponding to stimulus *p_j_*, let *pt_j, a_* correspond to the timepoint at which slice *j* was acquired exactly at the moment stimulus when stimulus *p_j_* was presented. In addition, let *pt_k, d_* correspond to the timepoint at which slice *k* was acquired precisely at timepoint *pt_j, a_* + Δ*t* (i.e., the end of the epoch). A given epoch *E_j_* then contains raw fMRI signal intensities as defined as the set of slices:

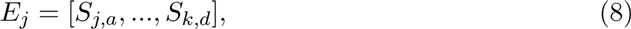

with the corresponding set of slice acquisition time points:

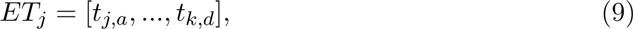

where each specific time point in this set is determined by Equation 3. Next, for each given epoch *E_j_* we compute the relative time difference *RET_j_* between the exact presentation time of the stimulus *pt*(*j, a*) and each time point in the epoch:

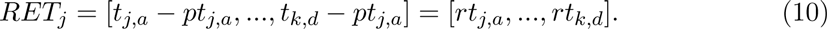

Importantly, note that because stimuli are presented precisely in-phase with the slice acquisition times, the relative times in *RET* are necessarily a subset of the actual slice acquisition times. This means that the total number of relative time points that is available in an epoch is equal to the number of slices included in the epoch. In turn, this means that the available temporal resolution in the epoch is equal to the slice sampling frequency.

Finally, we create a single epoch *L* with *r* whole-brain volumes

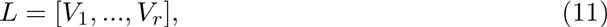

where *r* is determined by the ratio between the epoch length Δ*t* and the slice sampling frequency 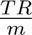. The corresponding matrix of volume acquisition times *LT* is determined by

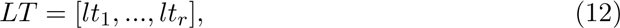

where each *lt* is determined by the function:

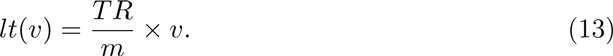

Each volume in the epoch *L* contains slices that are acquired at the same time point relative to the onset of the stimulus. This is achieved by combining slices from different epochs *E*_1_,…, *E_m_* on the basis of their *RET* values. Specifically slices 1,…, *m* can be combined into a whole-brain volume if their corresponding relative times *rt* match. For a given volume:

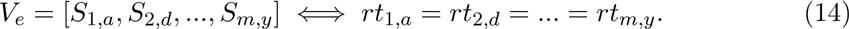

This then leads to an epoch of whole-brain volumes that do not contain any temporal distortions, and where volumes are available at a temporal resolution equal to the sampling frequency.

As a concrete example, consider the simplified situation represented in Figure 2. Here there are only three separate slices, slice1, slice2, and slice3 that provide whole-brain coverage. The TR is 3 s meaning that the sampling frequency is 1 s (see equation 3). Imagine there are three stimuli that are presented that are in-phase with the different slice acquisitions. Thus, for example, stimulus1 is presented at 0 s (in phase with slice1), stimulus2 at 7 s (in phase with slice2), and stimulus3 at 14 s (in phase with slice3). Then, during the presentation of stimulus1, slice1 will capture the state of the brain 0 s after the presentation of a stimulus, slice2 will capture the state of the brain 1 s after the presentation of a stimulus, slice3 2 s, and so on. During the presentation of stimulus2, slice2 will now capture the state of the brain 0 s after the presentation of the stimulus, slice3 1 s, and so on. After the presentation of all stimuli, each slice will have captured the state of the brain at all possible time points relative to the stimulus, and it is therefore possible to construct whole-brain volumes that are temporally correct and at a high temporal resolution.

### Statistical modeling of the BOLD signal

Statistical modeling of the fMRI BOLD signal differs fundamentally between the standard and the new Slice-Based framework. In the standard method, the fMRI BOLD signal is extracted from the data using a time-series approach in the context of a General Linear Model (GLM; Bandettini et al., 1993). A time-series approach entails that the fMRI data from an entire run is fitted to a single GLM model. In order to extract the time course of the BOLD signal, statistical methods have been developed that rely on so-called Finite Impulse Response (FIR) basis functions (e.g., Ollinger et al., 2001; Serences, 2004; Zarahn et al., 1997). In this type of analysis, each parameter in the GLM attempts to capture a different time point in the progression of the BOLD signal. This method requires choosing a certain epoch length, and then choosing a number of basis functions. Because in standard analyses the sample points are available at the resolution of the TR (see section above), the number of basis functions is usually chosen to be equal to the ratio of the epoch length and the TR. The GLM parameters of the FIR basis functions then capture the progression of the BOLD signal in the epoch at the temporal resolution of the TR. Within this method, the temporal resolution can be increased by adding additional sets of basis functions to the GLM and by accurately presenting stimuli out of phase with the TR. These additional parameters then attempt to capture additional timepoints in the progression of the BOLD signal (e.g., Dale, 1999; Josephs et al., 1997; Price et al., 1999; Toni et al., 1999). As has been repeatedly pointed out, a problem with this method is that increasing the number of parameters in the statistical model leads to a reduction of power (e.g., Lindquist et al., 2009).

By contrast, statistical modeling in the slice-based method does not rely on a time-series approach, but on a time point-by-time point approach. This approach to modeling data is common in recent approaches in electrophysiology (Janssen et al., 2014; Lage-Castellanos et al., 2010), but has not frequently been applied to fMRI data (but see Leung et al., 2000). This means that instead of fitting a single statistical model to a whole imaging run, separate statistical models are fitted at each individual time point in the epoch. To preserve sensitivity, we do not perform any data averaging, and analyses take place on all available data points from all stimuli at a given time point. A further aspect of these analyses refer to the particular choice of the baseline signal. In our analyses, we choose the baseline signal as those intensity values that are available at timepoint 1 in the epoch (i.e., the first time point). Statistical modeling then involves comparing intensity values at subsequent timepoints to those values obtained at baseline, and therefore leads to a data-driven extraction of the BOLD signal. Finally, in our analyses below we further refined this baseline procedure by including in the baseline not only the intensity values at timepoint 1 from a given voxel, but also the intensity values at timepoint 1 from this voxel across all slices. In other words, the baseline at a given voxel included intensities values at timepoint 1 from that voxel (i.e., x and y coordinates), but also those values at timepoint 1 from that same voxel (i.e., the same x and y coordinates) at all other slices (i.e., z coordinates). We hypothesized that this baseline procedure would include global signal information that is unspecific to any particular voxel, and therefore lead to a more accurate extraction of the BOLD signal across time. We will come back to this issue in the General Discussion.

### The current study

The goal of the current paper was to evaluate the temporal accuracy of BOLD signal extraction in the standard FIR method and Slice-Based methods. Specifically, we compared BOLD signal extraction using the FIR method without STC, the FIR method with STC, and the Slice-Based method. Note, however, that these comparisons are complex because they entail both a difference in the volume creation method and in the statistical modeling of the signal. That is, both FIR techniques rely on the standard method of whole-brain creation (i.e., time shifting of slices), and use a GLM time series technique to extract the BOLD signal. By contrast, the Slice-Based method uses a very different method of volume creation (i.e., combining slices collected at same relative timepoints), and uses a time point-by-time point technique to extract the BOLD signal. Here we did not intend to separate the contributions of each of these aspects of the procedures in our comparisons. However, as will become clear below, it will be rather obvious to identify whether limitations of each of the three techniques is due to the volume creation or statistical modeling method.

We evaluated the temporal accuracy of these three techniques in the context of simulated and real-world data. In Simulation 1, we evaluated the three methods’ abilities to recover the absolute timing of a simulated hemodynamic response from the fMRI data. Specifically, we simulated an fMRI imaging run in which a large patch of (fictitious) neural tissue was stimulated in a slow event-related design. Stimulus presentations generated a uniform hemodynamic response across the whole neural tissue. This large patch of neural tissue was covered by a set of adjacent fMRI slices that therefore each sampled the exact same hemodynamic response signal (with standard fMRI sampling parameters). We had two basic questions: First, how well does each method recover the ground-truth signal from the sampled BOLD data? Second, how well do the three methods extract the same BOLD signal on the set of adjacent slices? We evaluated these questions in terms of the basic Pearson correlation between the extracted BOLD signal and the expected ground-truth signal, and in terms of the Pearson interslice correlation of the extracted BOLD signal between the adjacent slices. If the Slice-Based method more accurately extracts the time course of the BOLD signal we expected a higher correlation with the ground-truth hemodynamic response, and a higher correlation of BOLD signals between adjacent slices compared to the standard methods.

In Simulation 2, we were interested in each method’s ability to recover the relative timing of two generated, temporally delayed, ground-truth hemodynamic responses. We evaluated each method’s ability to recover from the extracted BOLD signals the ground-truth temporal delays of the two hemodynamic responses. We conducted the same simulations for three different temporal delays between the two hemodynamic responses. We examined the extracted BOLD signal by the three methods in terms of its estimated Time To Peak (TTP) value, the time at Half the Maximum amplitude (HM) at the rising edge of the BOLD signal, and the mean correlation of the BOLD signals with their respective ground-truth hemodynamic responses. Again we reasoned that if the Slice-Based method more accurately extracts the time course of the BOLD signal, it would yield TTP, HM, and mean correlation values that more accurately reflect the ground-truth signals than those estimated by the standard methods.

Finally, we examined real-world fMRI data collected from 30 participants performing a slow event-related overt picture naming task. Previous research has revealed that the picture naming task shows robust activity across a large area in the primary motor cortex (e.g., Murtha et al., 1999; Price, 2012). We examined activity in three voxels across three adjacent slices in the left motor cortex that all showed strong involvement in the task. We hypothesized that these adjacent voxels would sample a very similar hemodynamic response. The question we had was therefore whether the three methods could extract a coherent BOLD signal across the three adjacent voxels in the left motor cortex. Specifically, we examined the interslice correlation between three BOLD signals from the adjacent slices, as well as the number of peaks in the BOLD signal across the three slices. We hypothesized that if the slice-based technique is less prone to temporal distortions, it would yield a higher interslice correlation as well as a smaller number of unique peaks than the standard methods.

## Methods

### Simulation 1 - Absolute timing of BOLD signal

100 simulations were performed in the software package R (v3.3.1). Each simulation took the form of an fMRI imaging run in which a large piece of (fictitious) neural tissue was repeatedly sampled at a number of different slice locations. Our simulation was set up such that this neural tissue would generate a hemodynamic response that was identical at each slice location. Each slice therefore samples the same hemodynamic response, and this leads to the expectation that the extracted BOLD signal of slice should be highly correlated. The question then is which of the three discussed techniques best recovers the absolute signal, and for which technique the signal is most similar across the three slices.

To simulate an fMRI imaging run, we generated a series of consecutive hemodynamic responses on the basis of a number of presented stimuli. Specifically, 60 stimuli presented at long 18 s intervals induced a series of hemodynamic responses that were modeled with a double gamma function with default parameters using the neuRosim package (v0.2-12; Welvaert et al., 2011). The precise onsets of the stimuli were constructed to be in-phase with the slice acquisition times determined by the fMRI parameters described below. The long interval between stimulus onsets meant that the hemodynamic response came back to baseline before the next stimulus onset. This long duration hemodynamic response signal was generated at a very high temporal resolution and represents the ground-truth hemodynamic response to each stimulus in our simulated fMRI imaging run. Finally, to make the simulation more probabilistic we added some Ricean noise with sigma=0.1 to the generated ground-truth hemodynamic response signal (Gudbjartsson & Patz, 1995).

This ground-truth hemodynamic signal was subsequently sampled in a standard fMRI fashion. For the first simulation the TR was 3.0 s, and there were 3 slices. On each slice there was only one voxel. We sampled each slice in a sequential fashion (1,2,3). This means that each voxel will obtain a sample every 3 seconds, and that adjacent voxels (on adjacent slices) will obtain a sample every 1 second. Note again that each slice samples the same underlying hemodynamic response signal because we assumed that the same hemodynamic response is present at all slice locations. This generated basic raw fMRI data with a BOLD signal time series at three voxels across three adjacent slices. The signal was then normalized to zero mean and unit standard deviation. This data formed the basis of all further analyses.

Volume creation took place in the two different ways described above. In the standard method volumes were created out of the three adjacent slices. It was assumed that each slice was acquired at the same point in time which was determined by the middle slice. This resulted in 362 volumes of three slices that were not slice time corrected. We then created a second set of 362 volumes to which we applied the FSL *slicetimer* function. This therefore led to a set of volumes that were slice time corrected. For the Slice-Based method, volumes were created that contained slices acquired at the same point relative to a stimulus. This resulted in a stimulus locked epoch of 18 volumes. Statistical extraction of the BOLD signal by the FIR and Slice-Based methods was performed on these data.

For the FIR methods, we constructed a design matrix with 6 basis functions (i.e., the 18 s epoch length divided by the TR of 3 s). To obtain a temporal resolution higher than the TR and equal to the resolution obtained using the Slice-Based method, two additional sets of 6 basis functions were added, yielding a total of 18 parameters in the design matrix. Each of these parameter sets corresponded to a set of onsets that coincided with a particular slice acquisition time. Each of the 18 basis functions attempted to capture a single timepoint in the progression of the BOLD signal. Specifically, parameter values were generated by a long sequence of zeroes and ones at various points in the timecourse of the BOLD signal since stimulus onset. This same design matrix was used for the FIR without STC and the FIR with STC methods. All statistical modeling was done using the linear modeling (*lm*) function of R.

For the Slice-Based method, the BOLD signal was extracted by comparing the signal between the baseline time point 1 and each subsequent time point in the epoched data. As mentioned earlier, the baseline was chosen as the values of time point 1 across all available slices. Modeling was performed using the same linear modeling function in R as in the previous methods.

Finally, we repeated this modeling exercise at different temporal resolutions. It has been argued that increasing the temporal resolution increases the number of parameters in the FIR based method, and this has a negative impact on the statistical power of the method (Lindquist et al., 2009). Here we were interested in seeing how increasing the temporal resolution impacted the power in the FIR and Slice-Based methods. We will define power here in terms of the maximum t-value detected in the time course of the BOLD signal. In our simulations the temporal resolution can be increased by keeping the TR equal while increasing the number of slices in the simulation, and by presenting the stimuli in-phase with these slice acquisitions. Specifically, we increased the number of slices to 6 and 12, and adjusted the stimulus onsets to coincide with the new slice acquisitions. This resulted in a TR/6 and TR/12 temporal resolution, respectively.

### Simulation 2 - Relative timing of BOLD signal

Simulation 2 also involved 100 simulations. This simulation took the form of an fMRI imaging run in which two (fictitious) pieces of neural tissue were repeatedly sampled. Our simulations were set up such that each neural tissue produced a neural response after the presentation of a stimulus. However, the hemodynamic response in the second neural tissue was slightly delayed relative to the hemodynamic response in the first neural tissue. Sampling of each piece of neural tissue was determined by assigning half the slices to one neural tissue, and the other half to the other piece of tissue. We systematically delayed the onset of the hemodynamic response in each neural tissue. This simulation was intended to ascertain to what extent each of the three aforementioned techniques is able to accurately recover the ground-truth temporal delay between the two generated hemodynamic responses.

In our first simulation we used 3 slices, meaning the first two slices covered the first piece of tissue, and the other slice the second piece of tissue. In this first simulation we systematically delayed the onset of the hemodynamic response in the second piece of neural tissue by 1 TR, 2/3 TR and 1/3 TR. To evaluate how extraction of the relative timing differences between three methods would function under different temporal resolutions, we increased the number of slices from 3 to 6 in a second set of simulations. All other aspects of Simulation 2 were identical to those of Simulation 1.

### Real-world data - Picture Naming

Finally, we attempted to evaluate the three methods in the context of real-world data. Although obviously we cannot know the ground-truth signal in these data, based on existing evidence we reasoned that picture naming should yield a similar hemodynamic response in a large portion of motor cortex (e.g., Murtha et al., 1999; Price, 2012). We then examined how well the three methods could extract coherent BOLD responses across three adjacent slices in each participant’s left motor cortex.

#### Participants

Thirty native speakers of Spanish took part in the experiment (20 females, 10 males, mean age around 22). Participants were students at the University of La Laguna, and received course credit or were paid 10 Euro. Twenty-nine participants were right-handed. The study was conducted in compliance with the declaration of Helsinki, and all participants provided informed consent in accordance with the protocol established by the Ethics Commission for Research of the university of La Laguna (Comité de Ética de la Investigación y Bienestar Animal).

#### Experimental setup and procedure

Two stimuli were used in the task: First, an image which participants were asked to name aloud, and second, a fixation cross (’+’) which indicated rest (see Figure 3 for an overview). Twenty-seven pictures were selected from an image database that contained standardized line-drawings that were normed on various aspects (Szekely et al., 2004). Only those images were selected that had names that were consistently produced across participants in the norming study (i.e., those with > 90% name-agreement).

**Figure 3:**
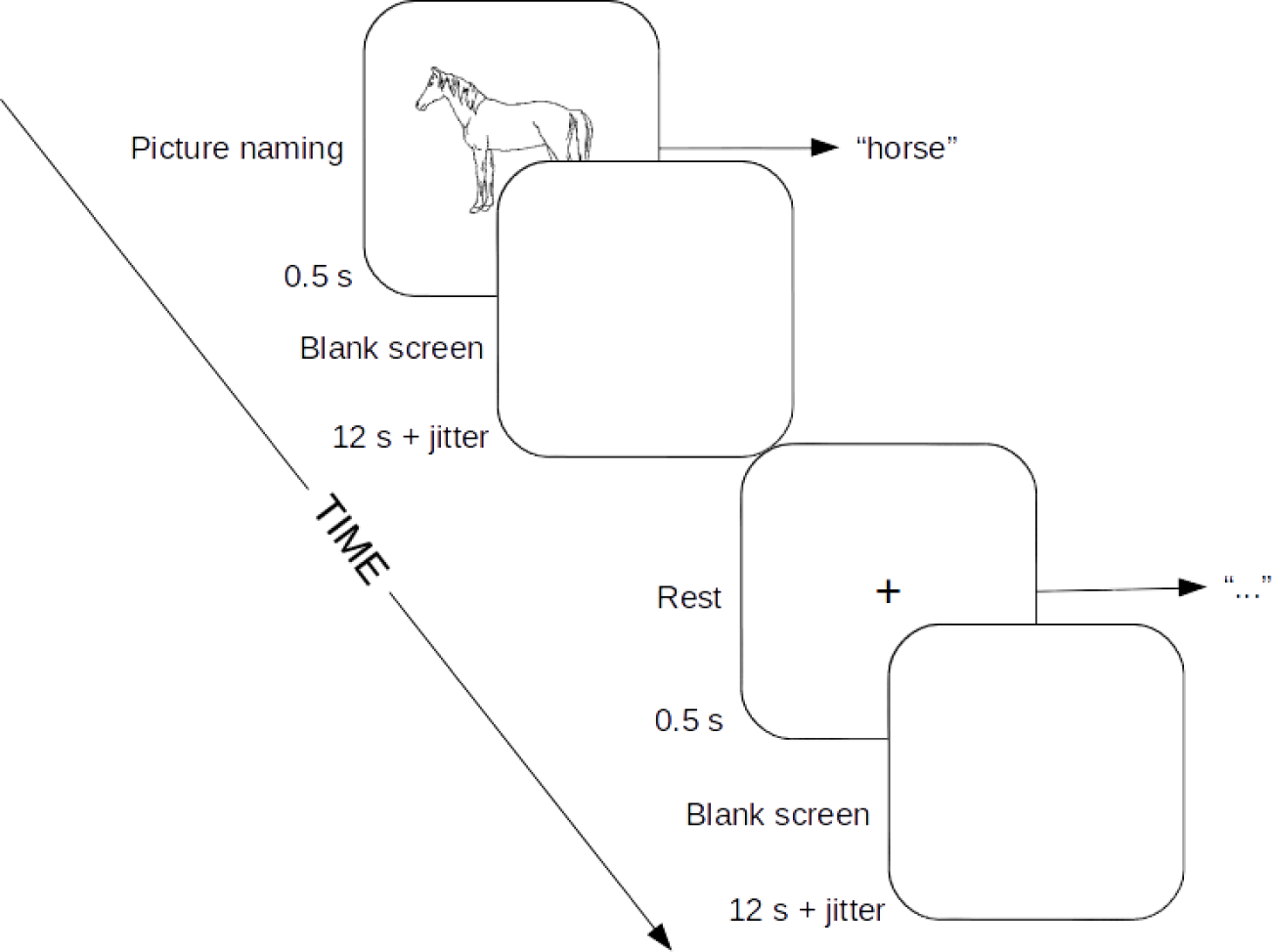
Temporal structure of the picture naming task used in the experiment. Stimuli consisted of either a picture or a fixation point that was presented for 0.5 s. Each stimulus presentation was followed by a blank screen that lasted for 12 s plus an additional jitter period. The jitter period was randomly selected without replacement from a uniform distribution of times that coincided with the slice acquisition times and ranged from 0 to 1855 ms in steps of 53 ms (see text for further details). Participants were instructed to name aloud presented pictures and remain quiet (i.e., rest) for presented fixation points. The order of stimuli presentation was fully randomized, and was different for every participant.

Stimuli were presented in a slow event-related design, where a stimulus was presented for 0.5 s followed by an ISI blank screen for 12 s plus an additional jitter period. The duration of the jitter period was randomly chosen without replacement from a uniform distribution of 36 times from 0 to 1855 ms in steps of 53 ms. This method of stimuli presentation complies with the requiremens of the slice-based method (see Figure 2 for further details). Stimulus presentation was directly synchronized with the MRI machine.

The Experiment involved three consecutive runs. In each run, 36 stimuli were presented, of which half were pictures and half were rest. In each run, nine different pictures were randomly selected and which were presented twice. Different pictures were selected for each run, and all twenty-seven pictures were presented in the experiment. For each run, the order of the stimuli was fully randomized on a by-participant basis. Stimulus presentation was controlled by Neurobs Presentations (v14). Participants in the scanner viewed the stimuli with MRI compatible goggles made by VisuaStim. These goggles provide an image resolution of 800 by 600 pixels at 60 Hz.

#### MRI acquisition parameters

MR-images were acquired using a 3T Signa Excite scanner (General Electric, Milwaukee, WI, USA) using a standard transmit/receive 8 channel gradient head coil. Head movement was strenuously avoided by fixating each participant’s head with spongepads inside the coil. T2*-weighted images were obtained using standard Gradient Echo, Echo Planar Imaging (EPI) sequences.

Each run started with 10 dummy volumes that allowed for steady-state tissue magnetization. Each volume contained 36 slices that were acquired top-down, axially and interleaved. Slice thickness was 3.7 mm with 0.3 mm gap. The FOV was 256 x 256 mm, matrix size 64 x 64, resulting in 4 x 4 x 4 mm isometric voxels. TR was 1908 ms, echo time (TE) 21.6 ms, and the flip angle 75^°^. This unusual TR was chosen because it was the fastest TR possible in the context of the other parameter settings and therefore would generate the maximum amount of data. In addition, 1908 is a multiple of 36 and this simplifies determining the slice acquisition times and stimulus presentation times. In each run 255 volumes were collected and lasted 8 minutes and 6 seconds.

Separate high resolution T1-weighted images were acquired using the 3D FSPGR sequence: TI/TR/TE: 650/6.8/1.4 ms, flip angle = 12^°^, 196 slices, slice thickness 1 mm, matrix 256 x 256, voxel size = 1 x 1 x 1 mm.

#### Pre-processing

Data pre-processing was minimal, and avoided any potential for temporal distortion of the signal (see also Discussion). First, spikes in the intensity values in the 4D datasets were removed using AFNI’s 3dDespike tool with default settings (Cox, 1996). Next, each 4D data set was motion corrected using FSL MCFLIRT, and low frequency drifts were removed with a high pass filter at 0.01 Hz (Smith et al., 2004). No smoothing was performed. Each data set was then slice time corrected using the FSL *slicetimer* function. Note that the slice time corrected data set was only used to extract data for the FIR with STC technique. Other techniques used the uncorrected data set.

#### BOLD signal extraction from left motor cortex

The minimally pre-processed datasets were then used to identify those voxels strongly activated during the picture naming task. We used the standard FSL FEAT procedure to detect activated voxels (Jenkinson et al., 2012). Precise picture naming onsets were extracted from the participant-specific Presentation log files. The rest periods were not explicitly modeled (Pernet, 2014). The expected HRF was modeled as a double gamma function with default parameters. We included the temporal derivative in the design matrix to ensure improved detection of signals that were slightly temporally delayed. Statistical modeling was performed in the context of the GLM. We only analyzed the first run of each participant.

We were specifically interested in three activated voxels on adjacent slices in each participant’s left motor cortex. To identify these three voxels we first created a mask of each participant’s left precentral gyrus using the lateralized Harvard-Oxford probabilistic atlas (Desikan et al., 2006). We manually removed any medial voxels included in the mask (e.g., SMA, pre-SMA) such that the mask only covered the lateral areas of the brain. Next we identified for each participant’s first run the voxel with the maximum t-value in the masked signal detection map, which corresponds to the voxel with the maximum t-value in the left motor cortex during the first run for a given participant. We then chose the voxel directly above (+z) and below (-z) this maximally activated voxel, resulting in thee voxels with the same x and y coordinates but differing z values (and therefore on adjacent slices). Finally, we extracted the raw BOLD time series from each participant’s motion corrected and temporally filtered data set at the three voxels identified by the procedure described above. This set of three vectors for every participant formed the input to the thee techniques described above.

Specifically, we again examined the BOLD signals extracted from these three voxels in left motor cortex using the FIR without STC, FIR with STC, and Slice-Based methods. The extraction was performed exactly as described above using the simulated data. We examined the mean interslice correlation, the mean number of unique peaks, the mean TTP, and the mean max t-value in the BOLD signal across the three slices for all participants. We extracted the BOLD signal in these thee voxels in left motor cortex at two temporal resolutions, the TR (1908 ms), and 1/2 TR (954 ms). We performed statistical comparisons of these values on a by-participant basis.

## Results

### Simulation 1

Qualitatively, as can been seen in Figure 4, it seems that the Slice-Based method (right most column) extracted the BOLD signals in closer correspondence with the ground-truth signal (dashed line in all figures) than both the FIR methods. For the FIR without STC method (left most column), the pattern is that the three slices appear to be phase shifted. This pattern is exactly what is expected under the standard volume creation method when three slices sample the same hemodynamic response with a 1 s delay (see Introduction). As also expected, this phase shift is reduced by the FIR with STC method (middle column). A final observation is that the FIR based methods seem to be detecting the BOLD signal with the same statistical significance as the Slice-Based method, suggesting that the FIR based statistical modeling method is working well, and that the problem in terms of temporal accuracy lies with the volume creation method.

**Figure 4:**
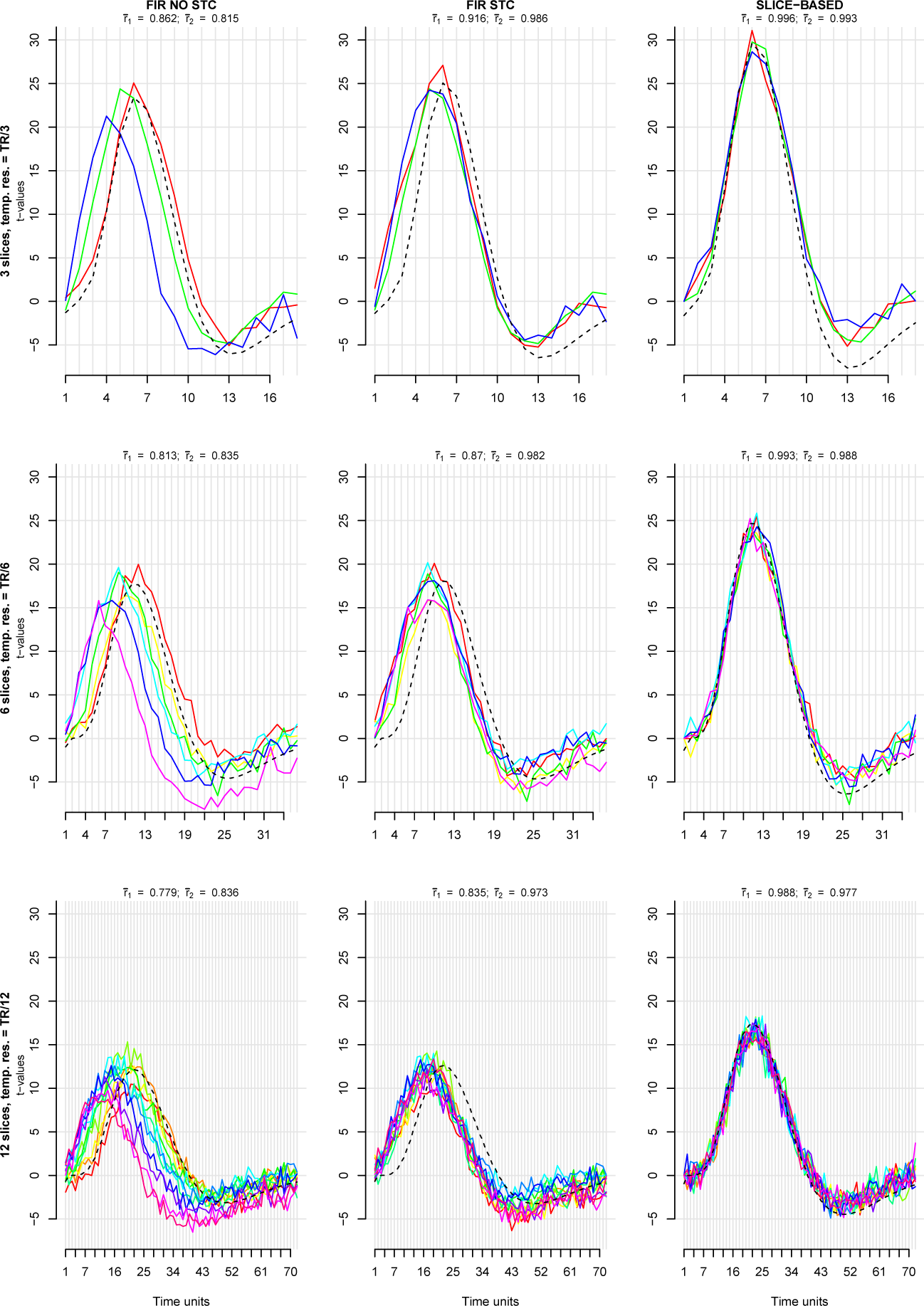
Detection of the absolute timing of a single ground-truth signal (black dotted line). The same signal is sampled by several adjacent slices (colored lines). Examples from signal extraction using the FIR method without STC (left column), with STC (middle column) and the Slice-Based method (right column) at a temporal resolution of the TR divided by 3 (top-row), by 6 (middle row), and by 12 (bottom row). X-axis represent time in arbitrary units, and y-axis represents t-values obtained from model fitting using the FIR and slice-based methods. Figure titles list the mean correlation between the ground-truth signal and the signal from each slice 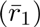, and the mean interslice correlation 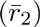 across all simulations. Each row contains data from a single simulation. Note how the slice-based method more accurately extracts the absolute timing of the BOLD signal (see text for further details).

These qualitative impressions were further confirmed in quantitative analyses of the simulation data. At the lowest temporal resolution (top row Figure 4), the correlation between the ground-truth signal and the mean detected signal across all slices was higher for the Slice-Based method 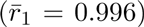, than for the FIR without STC 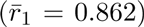, and FIR with STC 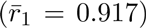, suggesting an improved accuracy of around 16% for the Slice-Based method relative to the FIR without STC method. Note that we do not report any detailed statistics here due to the largely deterministic nature of the simulations which led to all reported numerical differences here to be highly statistically significant. In addition, the Slice-Based method revealed a more similar extracted BOLD signal across the three slices. The mean interslice correction for the Slice-Based method was significantly higher 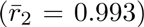 than for the FIR without STC 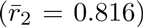, and FIR with STC 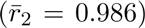, suggesting a 21% improvement in accuracy for the Slice-Based method. Note that this improved accuracy was preserved with increasing temporal resolutions (the rows of Figure 4). Specifically, the correlation with the ground-truth signal increased by 22% and 27% for the TR/6 and TR/12 temporal resolutions in the Slice-Based method. Similarly, the interslice correlation increased by 18% and 17% for the TR/6 and TR/12 temporal resolutions for the Slice-Based method.

### Simulation 2

A visual impression of Figures 5 and 6 suggested that the Slice-Based method was also superior in detecting the relative timing of the two ground-truth hemodynamic responses (dashed lines). This seemed to be the case independently of their temporal delay, and independently of the temporal resolution. For example, although the FIR without STC technique (top left, Figure 5) seemed to extract a hemodynamic response for the slices 1 and 2 (red and green lines) that seemed to arise earlier than the second hemodynamic response for slice 3 (blue), neither the absolute nor the relative timing of these two hemodynamic responses seemed to be correct, and the relative timing seemed increasingly incorrect with increasing temporal resolution (top left Figure 6). By contrast, the Slice-Based technique seemed to accurately capture both the absolute and relative timing.

**Figure 5:**
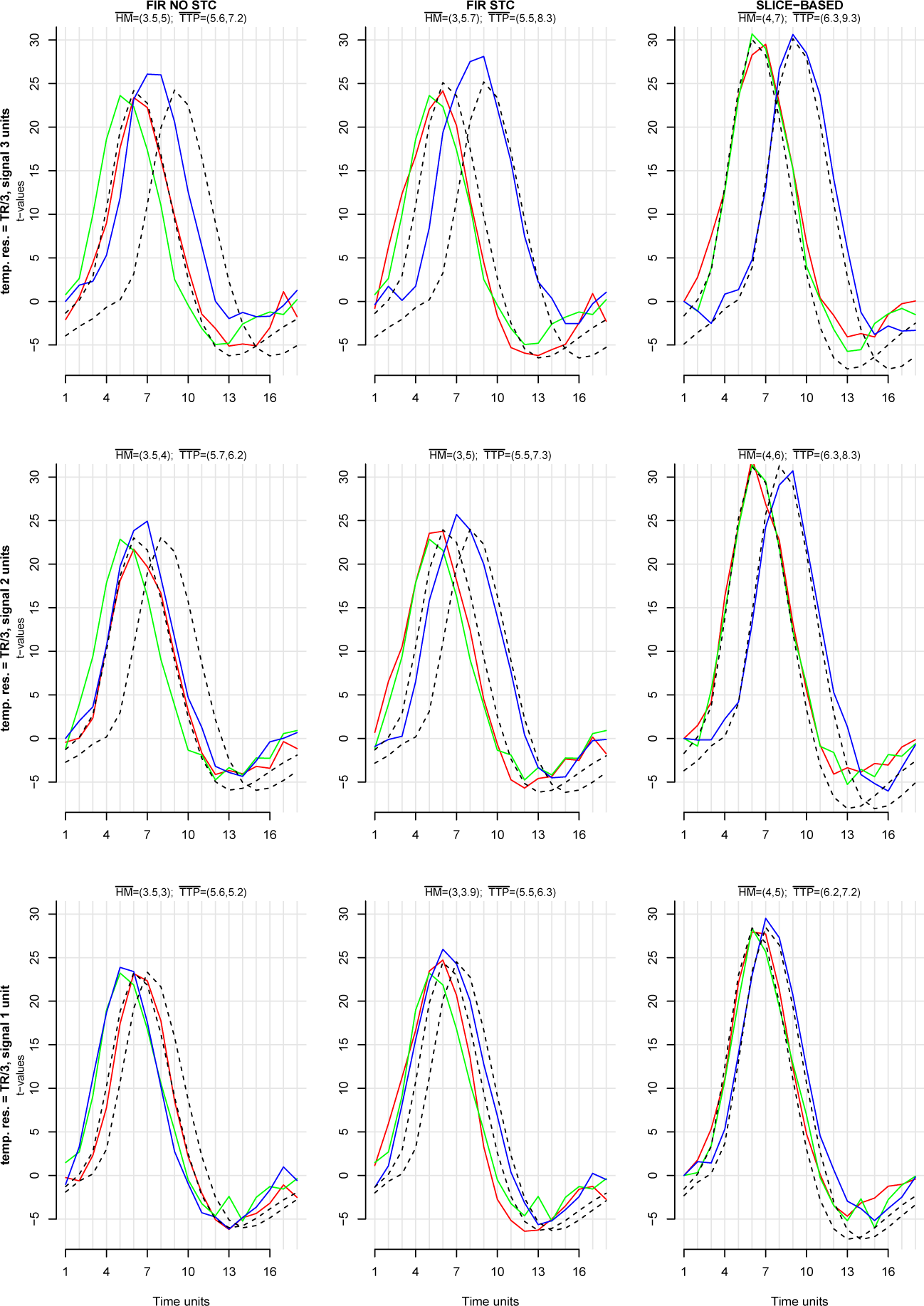
Detection of the relative timing of two ground-truth signals (black dotted lines) whose relative onsets are delayed by 1 TR (top-row), 2/3 TR (middle row), or 1/3 TR (bottom row). Examples from signal extraction using the FIR method without STC (left column), with STC (middle column) and the Slice-Based method (right column) at a fixed temporal resolution of the TR divided by 3. Each row contains data from a single simulation. Figure titles list the mean timepoint at which the curve reaches Half of the Maximum amplitude (HM), the mean Time To Peak (TTP), and the mean correlation of the extracted BOLD signal for the first and second ground-truth signal 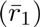 across all simulations. Note how the slice-based method more accurately extracts the relative timing of the two BOLD signals.

**Figure 6:**
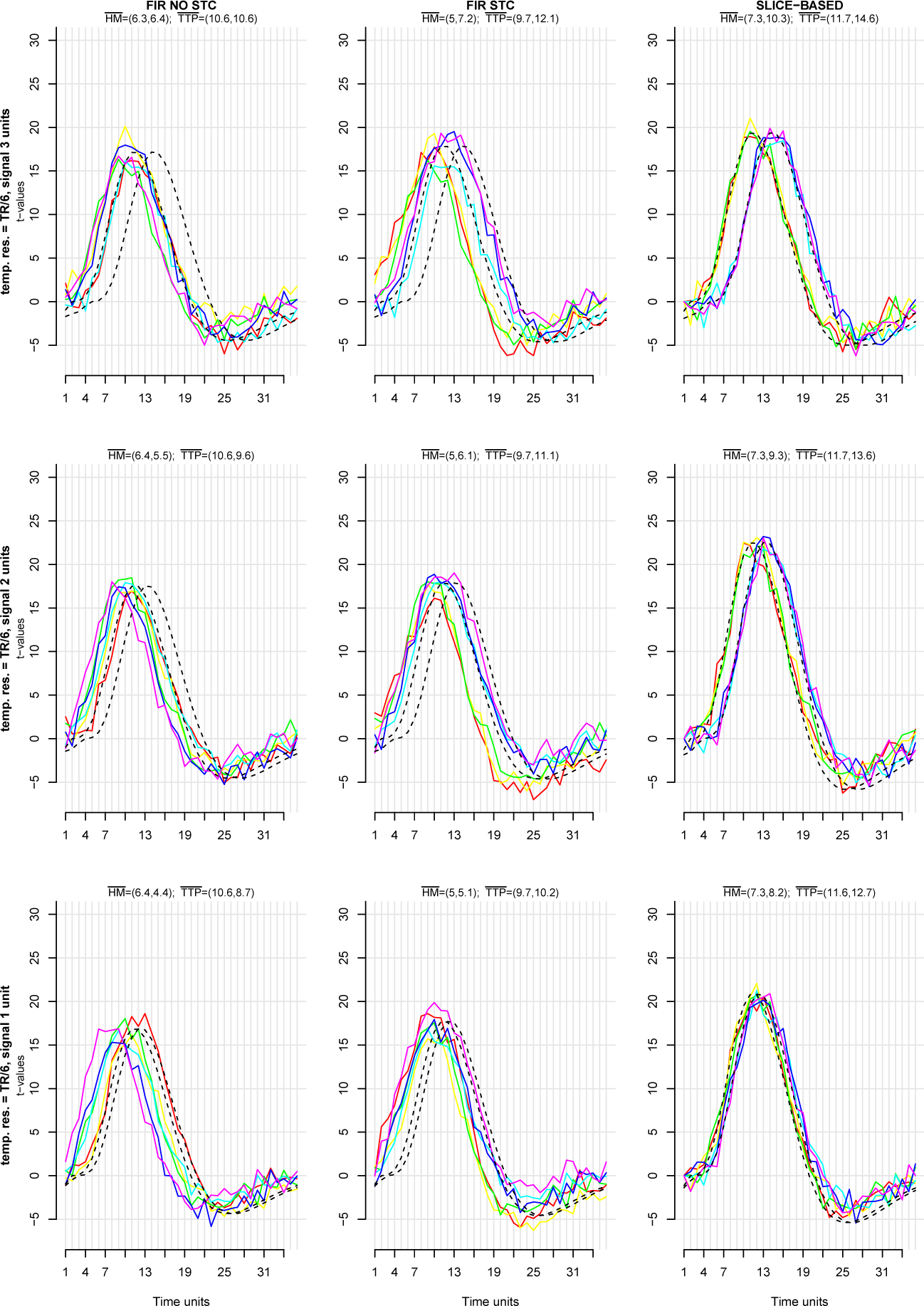
Detection of the relative timing of two asynchronized ground-truth signals (black dotted lines) across several slices (colored lines) at a fixed temporal resolution of TR/6. Each row contains data from a single simulation. Again note how the slice-based method more accurately detects the relative timing of the two BOLD signals.

Quantitatively, at a temporal resolution of TR/3 and a ground-truth temporal delay of 3 time units (see top row Figure 5), the Slice-Based method detected a mean difference of the HM value between the first and the second BOLD signal of exactly 3 (4 for the first, 7 for the second signal), and 3.1 for TTP (6.2 for the first, 9.3 for the second signal). The mean correlation with the first ground-truth signal was 1, while the mean correlation with the second ground-truth signal was 0.99, suggesting a mean accuracy of 99.5%. The FIR without STC method detected a mean difference of the HM value of 1.5, and a mean difference of 1.6 for the TTP value. The correlation with the mean ground truth signal was 0.865, suggesting a 86.5% accuracy. Finally, the FIR with STC method detected a mean difference of the HM value of 2.8, and a mean difference of 2.8 for the TTP value. The correlation with the mean ground truth signal was 0.918, suggesting a 91.8% accuracy.

These accuracy values remained constant for decreasing temporal delays. At 2 and 1 units delay (middle and bottom rows of Figure 5), the Slice-Based method was 99.5% accurate, while the FIR without STC and the FIR with STC were 86.6% and 91.2% accurate, respectively.

However, the accuracy values for the FIR based methods severely changed with increasing temporal resolution caused by a poor correlation with the second ground-truth signal. At a temporal resolution of TR/6 and a temporal delay of 3 units (see top row Figure 6), accuracy for the FIR without STC dropped to 57.9%, and to 67.1% for the FIR with STC. The Slice-Based method was 90.6% accurate. At 2 temporal units delay, the FIR without STC method slightly increased to 66.2% accurate, the FIR with STC method was 74.9% accurate, and the Slice-Based method was 95.7% accurate. Finally, at 1 unit temporal delay, the FIR without STC was 66.2% accurate, the FIR with STC was 81.8% accurate, and the Slice-Based method was 98.7% accurate.

### Picture Naming data

Figure 7 provides an overview of the extracted BOLD signals for the three techniques in the three voxels in left motor cortex for three representative participants from a single imaging run with 18 picture naming stimuli. The visual impression of this figure is that the Slice-Based method yielded BOLD signals across adjacent slices that are more similar than those of the other two methods. In addition, it appeared that the BOLD signals extracted by the Slice-Based method had fewer unique peaks than those of the other methods. Figure 8 relied on the same data but BOLD signals are now extracted at twice the temporal resolution (0.5 TR versus 1 TR). Here we saw a similar pattern where the Slice-Based method yielded smoother signals that appeared similar across the three slices. A final observation was that the maximum t-values appeared lower for the Slice-Based method than for the other methods.

**Figure 7:**
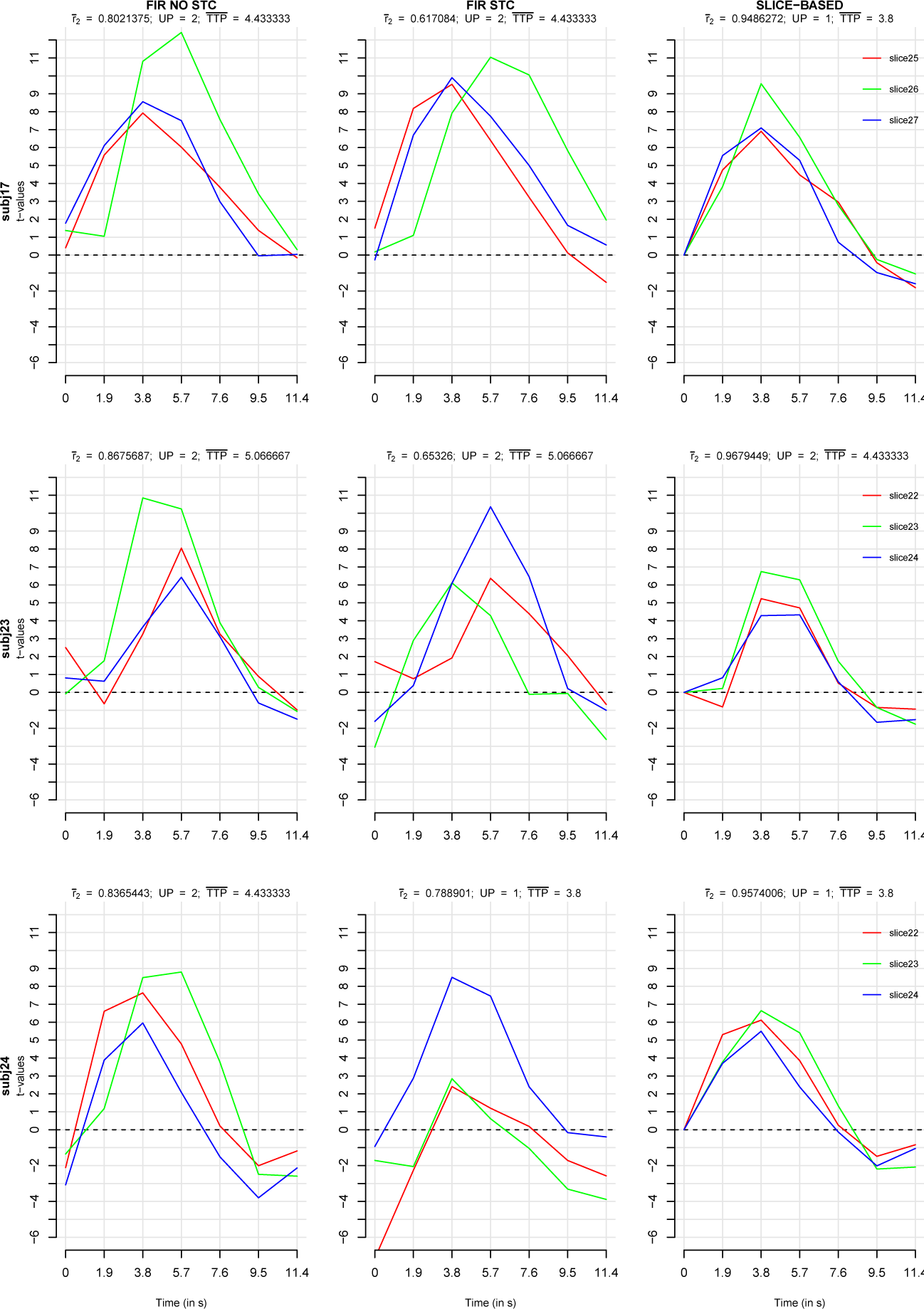
Method comparison using real data from left motor cortex activity obtained using the picture naming task. BOLD signal extracted using the FIR method without Slice Time Correction (left column), with Slice Time Correction (middle column) and the Slice-Based method (right column) at a fixed temporal resolution of the TR (1908 ms). Signals are extracted from three voxels that appear on adjacent slices (see legend) in the left motor cortex in three representative subjects (top, middle, and bottom row for subjects 17, 23, and 24, respectively). Figure titles list the interslice correlation 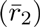, the mean number of Unique Peaks (UP), and the mean Time To Peak (TTP) for the extracted signals in the graph. Note how the slice-based method yields more temporally coherent signals from adjacent slices.

**Figure 8:**
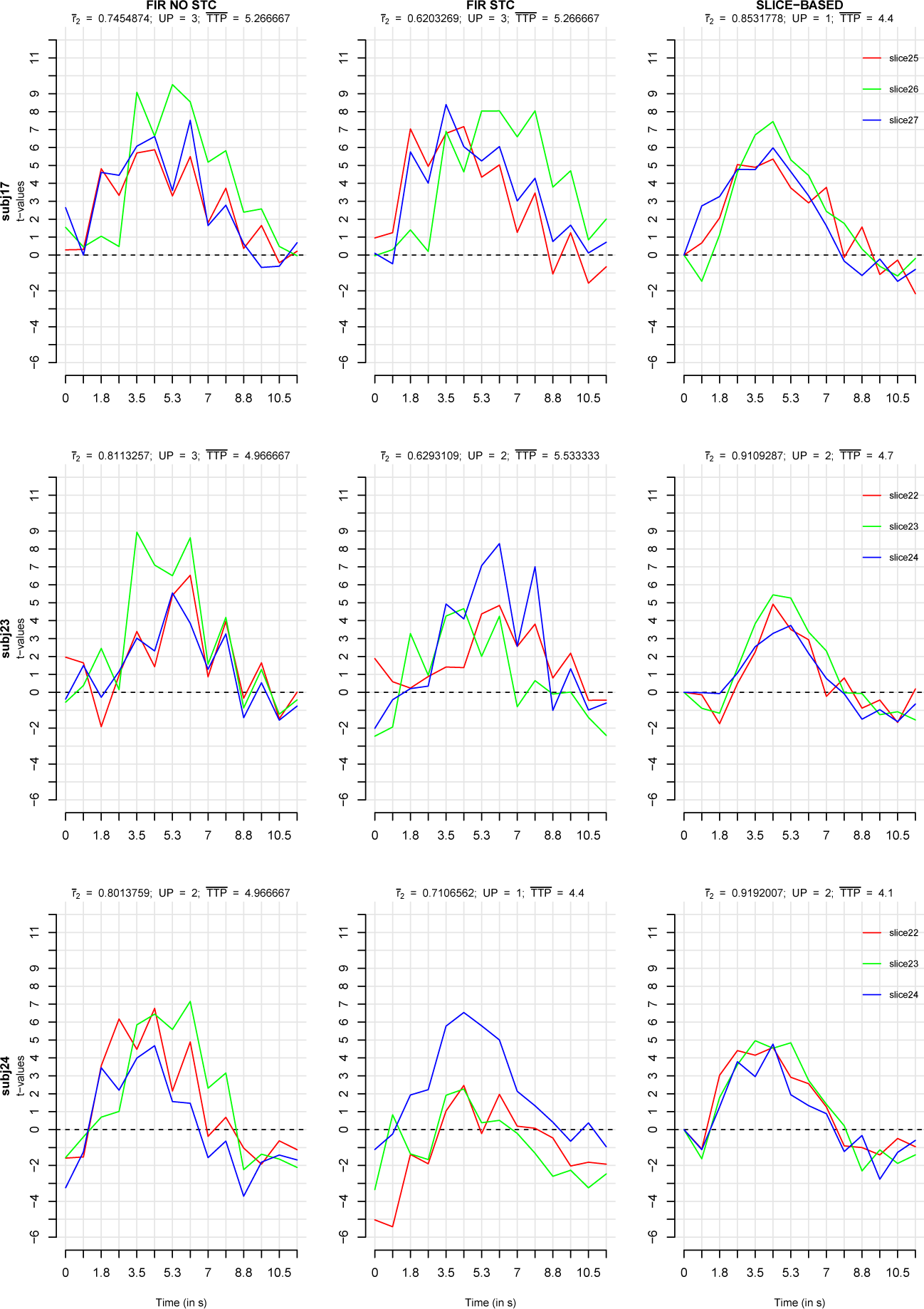
Method comparison using real data from left motor cortex activity obtained using the picture naming task. BOLD signal extracted using the FIR method without Slice Time Correction (left column), with Slice Time Correction (middle column) and the Slice-Based method (right column) at a fixed temporal resolution of 1/2 TR (954 ms). Other aspects identical to those used to obtain Figure 7. Note how despite an obvious drop in statistical power due to the reduced number of data points available per time point, the slice-based method yields more temporally coherent signals from adjacent slices.

As can be seen in Figures 9 and 10, these visual impressions were largely confirmed by statistical comparisons. For the standard TR resolution (see Figures 7 and 9, panel A), the mean interslice correlation across all 30 participants differed between the Slice-Based method and the FIR without STC method (*t*(29) = 4.65, *p* < 0. 001), suggesting higher mean interslice correlations in the Slice-Based method (0.83 vs 0.69). In addition, the interslice correlation for Slice-Based method differed from the FIR with STC method (*t*(29) = 6.63, *p* < 0. 001), indicating higher interslice correlations in the Slice-Based method (0.83 vs 0.35). Finally, the interslice correlation differed between the FIR without STC and FIR with STC (*t*(29) = 5.00, *p* < 0. 001), revealing a higher interslice correlation in FIR without STC than with STC (0.69 vs 0.35).

**Figure 9:**
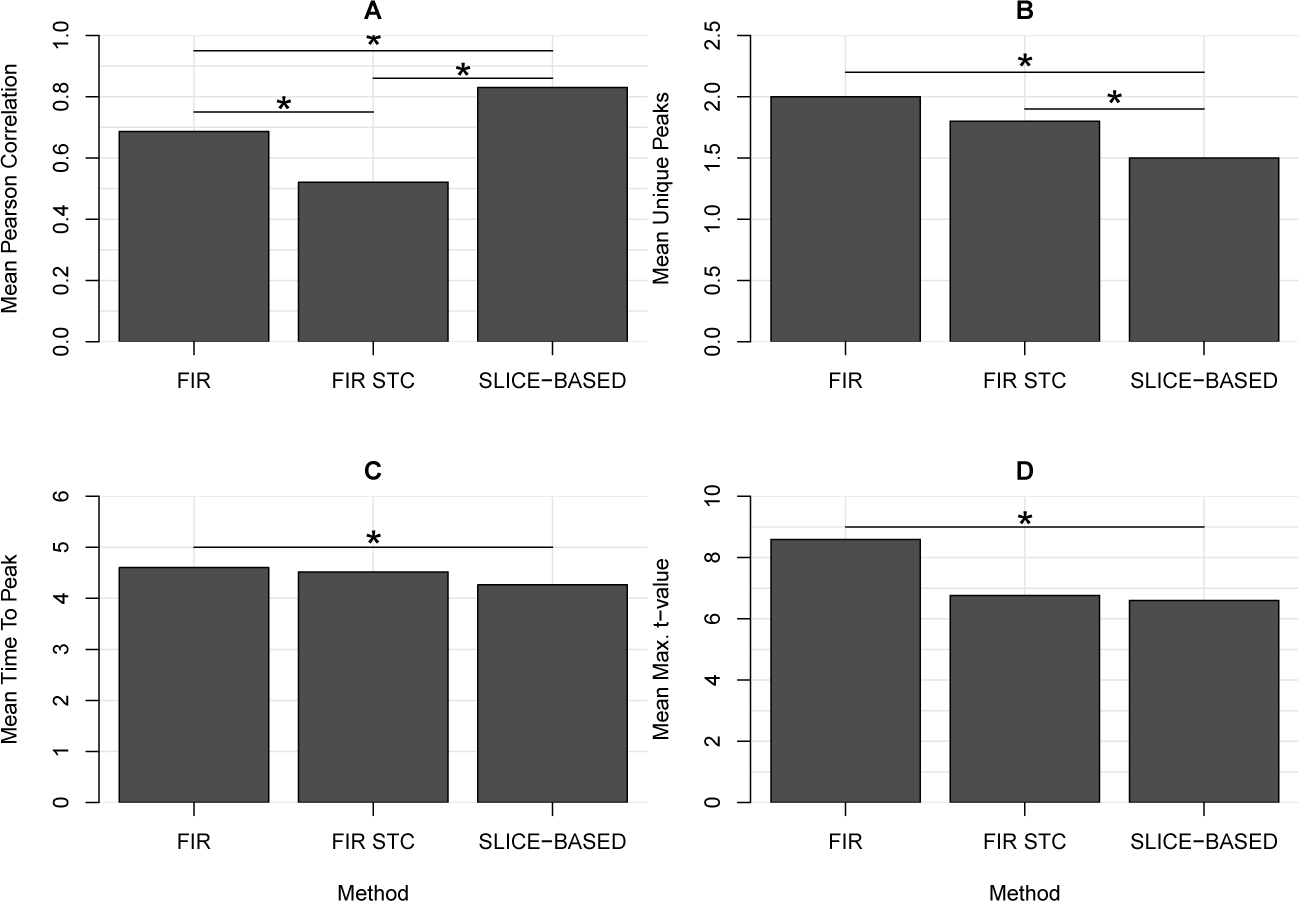
Graphical presentation of the overall relationship of the extracted BOLD signals from three adjacent slices covering the left motor cortex. The relationship was assessed using the mean inter slice Pearson Correlation (A), mean number of Unique Peaks (B), mean Time To Peak (C), and mean Max t-value (D) for the FIR, FIR with Slice Time Correction, and Slice-Based methods. Mean values obtained from 30 participants performing the picture naming task at TR resolution (1908 ms). The slice-based method yields improved inter-slice correspondence, suggesting improved temporal accuracy. See text for details.

**Figure 10:**
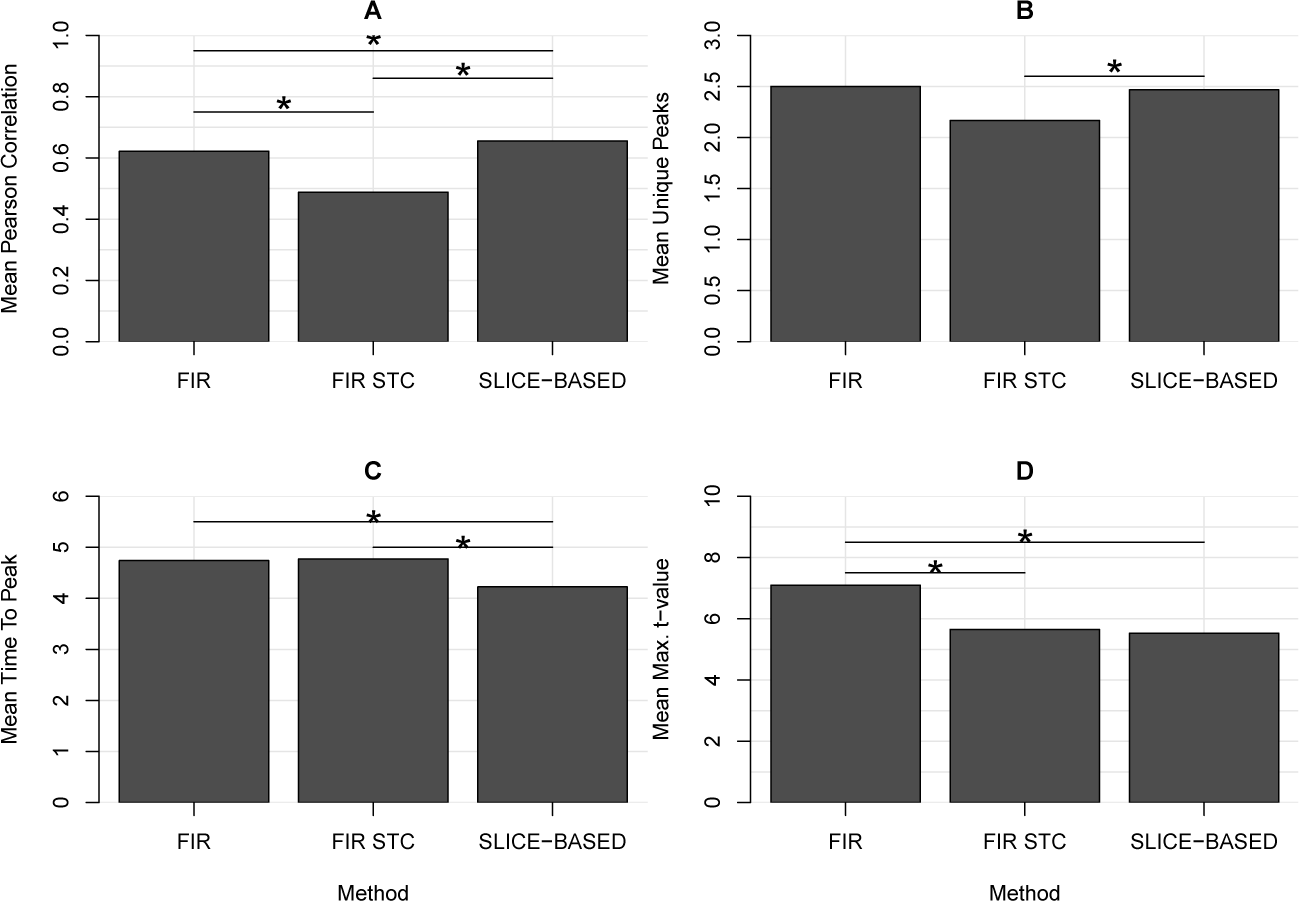
Graphical presentation of the overall relationship of the extracted BOLD signals from three adjacent slices covering the left motor cortex. The relationship was assessed using the mean inter slice Pearson Correlation (A), mean number of Unique Peaks (B), mean Time To Peak (C), and mean Max t-value (D) for the FIR, FIR with Slice Time Correction, and Slice-Based methods. Mean values obtained from 30 participants performing the picture naming task at 0.5 TR resolution (954 ms). The slice-based method yields improved inter-slice correspondence, suggesting improved temporal accuracy. See text for details.

Similarly, the mean number of unique peaks (see Figures 7 and 9, panel B) differed between the Slice-Based method and the FIR without STC method (*t*(29) = 4.00, *p* < 0.001), suggesting a lower mean number of unique peaks for the Slice-Based compared to the FIR without STC method (1.5 vs 2 peaks). Furthermore, the mean number of peaks differed between the Slice-Based method and the FIR with STC method (*t*(29) = 5.75, *p* < 0.001), indicating a lower number of mean peaks for the Slice-Based than the FIR with STC method (1.5 vs 2.3). Finally, the mean number of peaks differed between the FIR with and without STC (*t*(29) = 2.34, *p* < 0.03), yielding a lower mean number of peaks in the FIR without STC method (2 vs 2.3).

The mean TTP value (see Figures 7 and 9, panel C) also differed between Slice-Based and the FIR with STC method (*t*(29) = 2.81, *p* < 0.009), suggesting an earlier TTP for the Slice-Based compared to the FIR with STC method (4.28 vs 4.60 s). Other comparisons did not reach significance.

Finally, the mean maximum t-value in the time course of the BOLD signal (see Figures 7 and 9, panel D) differed between the Slice-Based and the FIR without STC methods (*t*(29) = 6.83, *p* < 0.001), suggesting a higher t-value for the FIR without STC than the Slice-Based method (6.60 vs 8.60). Similarly, the FIR without STC differed from the FIR with STC method (*t*(29) = 4.78, *p* < 0.001), indicating a higher t-value for the FIR without STC than for the FIR with STC method (8.60 vs 6.76). Other comparisons did not reach significance.

A similar pattern of results was found for the high temporal resolution data (see Figures 8 and 10). The mean interslice correlations (see Figures 8 and 10, panel A) differed between the Slice-Based and the FIR with STC method (*t*(29) = 5.66, *p* < 0.001), indicating a higher mean interslice correlation for the Slice-Based method (0.66 vs 0.34). Similarly, the correlation differed between FIR without STC and the FIR with STC methods (*t*(29) = 4.75, *p* < 0.001), pointing to a higher mean interslice correlation for the FIR without STC method (0.62 vs 0.34).

The mean TTP (see Figures 8 and 10, panel C) differed between the Slice-Based and the FIR without STC method (*t*(29) = 3.55, *p* < 0.002), indicating again an earlier TTP for Slice-Based method (4.2 vs 4.74). Other comparisons did not reach significance.

Finally, the mean maximum t-value (see Figures 8 and 10, panel D) differed between the Slice-Based and the FIR without STC method (*t*(29) = 6.10, *p* < 0.001), suggesting higher maximum t-values for the FIR without STC method (7.09 vs 5.54). In addition, the maximum t-value differed between FIR with and without STC (*t*(29) = 4.16, *p* < 0.003), indicating a higher t-value for the FIR without STC (7.09 vs 5.75).

## Discussion

In the current study we evaluated a new framework for fMRI data analysis called Slice-Based fMRI. The main purpose of the Slice-Based framework is to improve the temporal accuracy by which the fMRI BOLD signal can be extracted from the data. The framework relies on two core principles. First, it relies on a new method of volume creation. Specifically, instead of the standard method of time-shifting slices into whole-brain volumes, the Slice-Based method creates volumes in which all slices are acquired at the same time point relative to a presented stimulus (see Figures 1 and 2). Second, the method relies on a non-standard statistical modeling approach to BOLD signal extraction. In particular, whereas the standard method relies on a time-series approach in which a single GLM is fitted to the data from an entire imaging run, the Slice-Based method relies on a time point-by-time point approach in which separate statistical models are fitted at each time point in epoched stimulus locked data. We have used simulated data to compare the performance of the Slice-Based and standard methods in terms of their abilities to recover the absolute timing of a ground-truth signal, and in terms of their abilities to recover the relative time course properties of two temporally delayed ground-truth signals. Real-world data was used to assess the expected similarity of BOLD signals on three adjacent slices in left motor cortex during a language production task with 30 participants. Our results provide evidence that the Slice-Based method improves the temporal accuracy by which the BOLD signal can be extracted from the data. In this discussion section we touch upon a number of relevant issues.

### Impact of volume creation method

In Simulation 1, we tested the situation in which a large patch of (fictitious) neural tissue generated a series of uniform ground-truth hemodynamic responses when exposed to a series of consecutively presented stimuli in a slow event-related design. This large patch of tissue was covered by a number of slices that sampled the hemodynamic response in a standard sequential fMRI fashion. We compared the Slice-Based technique against two standard methods for BOLD signal extraction: FIR without STC and FIR with STC. In this first simulation, we were interested in two issues: First, how well do the standard and Slice-Based methods extract the absolute time course of the hemodynamic response, and second, is the BOLD signal extracted between the slices similar as would be expected? The data revealed that the FIR without STC method detected the ground truth signal with approximately 82% accuracy, and the FIR with STC with approximately 87% accuracy. The Slice-Based method detected the ground-truth signal with approximately 99% accuracy. In addition, the interslice correlation for the FIR without STC was 83%, and for the FIR with STC 98%. The Slice-Based method detected the interslice correlation with 99% accuracy.

How can these differences in temporal accuracy between the three methods be explained? Consider, for example, the findings displayed in the top-row of Figure 4. Here only three slices were used and the estimated ground-truth signal is represented by the dashed line. In the FIR without STC method (top left), the BOLD signals extracted across the three slices appear shifted in time, where the first slice (red line) appears later than the second slice (green line) which in turn appears later than the third slice (blue line). This state of affairs can be explained by the properties of the standard volume creation method. Recall that in this method, volumes are created by time shifting slices to a given reference slice. This means that at any time point in this epoch, slices within a volume were not acquired at the same moment in time. Thus, at time point 1, data for slice 1 was acquired 1 s earlier than that of reference slice 2, and data for slice 3 was acquired 1 s later than that of reference slice 2. The same is true for the other time points. These temporal inaccuracies reveal themselves as phase shifts of the BOLD signal for the respective slices in the epoch. This explains the temporal inaccuracy of the FIR without STC method.

The FIR with STC method is designed to alleviate these phase shifts caused by the volume creation method (Henson et al., 1999; Sladky et al., 2011). This method works by reversing the phase shift of the signals to become more in line with an arbitrarily chosen reference slice. In the FSL *slicetimer* function, the reference slice is by default the middle slice in the series. The effect of the STC function can be clearly seen by comparing the signals from the FIR without STC (left column) to those extracted in the FIR with STC method (middle column). Specifically, it can been seen that whereas the green line (middle slice, reference) is not adjusted, the red and blue signals are adjusted to become more like the green reference signal. Note that here the middle slice was a good choice for a reference slice because it increased the mean correlation with the ground-truth signal. However, it should be clear that this choice cannot be a-priori known, and hence, it is not obvious that STC will necessarily lead to a better detection of the actual ground-truth signal. If for example, the third slice (blue line) were chosen as a reference slice, the mean correlation would have decreased.

Thus, signal detection in the standard method is hampered by the volume creation method, and may be further hampered by the STC function. No such problems arise in the Slice-Based method. This is because in the Slice-Based method, each volume in the epoch contained slices that were all sampled at the same moment in time relative a presented stimulus. Thus, at time point 1 in the epoch of the Slice-Based method (top right), data for slice 1 was acquired at exactly the same moment in time relative to the presentation of the stimulus as the data for slice 2 and slice 3. This method therefore avoids phase shifts in the signal among the slices, and leads to highly consistent signals across the three adjacent slices. This particular method of volume creation also means that no STC is necessary for the Slice-Based data. In turn this leads to a highly accurate detection of the absolute time course of the signal, and to a high degree of interslice correlation. This is the case even under conditions of reduced power such as when the temporal resolution is increased (middle and bottom row Figure 4).

In Simulation 2, we tested the situation in which two (fictitious) neural patches generated two ground-truth hemodynamic responses. As before, the hemodynamic responses were generated by a series of presented stimuli, simulating a slow event-related fMRI imaging run. Crucially, these two hemodynamic responses were systematically temporally delayed. The two neural tissues were sampled by a set of adjacent slices. Our question was whether the standard and Slice-Based methods were able to recover the ground-truth temporal delays from the sampled BOLD signals. The data revealed that the FIR without STC method detected the ground truth temporal delays with approximately 74% accuracy, and the FIR without STC with approximately 83% accuracy. The Slice-Based detected the ground-truth temporal delays with approximately 97% accuracy.

The reason why the standard methods do not work well in Simulation 2 is the same as for Simulation 1. The volumes at each time point in the epoch contain slices that were not all acquired at the same moment in time (see Figure 5). As explained above, this introduces phase shifts in the sampled signal and produces temporal distortions. In turn this leads to a poor recovery of the ground-truth temporal delays in the standard FIR without STC and with STC. By contrast, given that in the Slice-Based method each volume in the epoch contains slices that were sampled at the same moment in time relative to a presented stimulus, no temporal distortions occur. In turn this leads to the accurate detection of the ground-truth temporal delays, and to a high mean correlation with the time course of the two absolute signals. Note that both simulations were tested under conditions of increasing temporal resolution, and that the Slice-Based maintained high accuracy under such conditions.

Finally, we evaluated the standard and Slice-Based methods using real-world data (see Figure 7). Specifically, we obtained fMRI data from 30 participants in a picture naming task. Previous research has demonstrated that picture naming strongly relies on activity in motor cortex (Murtha et al., 1999; Price, 2012). Assuming a similar hemodynamic response in the portion of left motor cortex covered by three adjacent slices, we asked the question of whether the three different techniques would be able to detect consistent neural activity in the three adjacent slices. Our data revealed that the interslice correlation was significantly higher in the Slice-Based method compared to the other two methods. In addition, the mean number of unique peaks in the signal across the three slices was consistently lower for the Slice-Based method than for the standard method. We found this result with a temporal resolution of the TR and with an increased temporal resolution at 0.5 TR. This therefore suggests that the Slice-Based method also extracts more consistent BOLD signals using real-world data.

The data presented in this report therefore reveal the limitations of the standard method of volume creation and the required STC function. Combining the accuracy values across all simulations, the FIR without STC detected the ground-truth signal with mean accuracy of 76% (0.24 sd), and the FIR with STC 86% (sd 0.14). By contrast, the Slice-Based method detected the ground-truth signals with an accuracy of 98% (sd 0.05). Thus, these data reveal that the Slice-Based method is about 22% more accurate, and an order of magnitude more precise (smaller variance). For researchers interested in extracting temporally accurate BOLD signals, the Slice-Based method offers a promising alternative. We will discuss future applications of the method further below.

### Impact of statistical method

The Slice-Based method not only differs from the standard method in terms of how volumes are created, but also in terms of how the statistical modeling of the fMRI data takes place. Were these two aspects confounded in the current results? We believe not, given that the FIR basis functions of the standard method extracted the BOLD signals with high statistical confidence. This means that the standard statistical methods revealed a correct representation of the BOLD signal, and that the temporal inaccuracies were introduced by the volume creation method. In other words, our results cannot be taken as evidence that the method of statistically extracting the BOLD signal using FIR basis functions was the reason for the temporal inaccuracy. The temporal accuracy in the fMRI data is primarily determined by the volume creation method.

That being said, the statistical methods of extracting the BOLD signal did differ substantially between the two methods. It is therefore worthwhile to further explain these differences, and their implications for the interpretation of the results. Specifically, we will argue that whereas the standard method yields effects of variables that reflect correlations between the stimulus presentation scheme and the observed data, the Slice-Based approach yields effects of variables that directly reflect differences in the fRMI signal intensity values. These differences have important consequences for the interpretation of the results.

As mentioned in the Introduction, in the standard method, BOLD signal extraction relies on a time-series approach where data is fitted with a GLM (e.g., Bandettini et al., 1993; Friston et al., 1994; Lindquist et al., 2009). In the design matrix of the GLM, independent variables such as the FIR basis functions used in the current study represent the time *when* a particular stimulus is present and absent (Dale, 1999; Josephs et al., 1997; Serences, 2004). These time-varying variables are then fitted against the BOLD fluctuations measured at each voxel. Effects of such variables therefore represent the degree to which the stimulus presentation scheme (present and absent) correlated with the observed data across time. For example, in the context of the current picture naming experiment, a high t-value in a given voxel reflects a high degree of (partial) correlation between the temporal patterning of turning on and off the picture stimulus and the temporal increases and decreases in the observed BOLD signal intensities.

By contrast, in the time point-by-time point approach used by the Slice-Based framework, independent variables represent Time with generally two levels (e.g., baseline vs another timepoint in the epoch). An effect of a variable in this type of model represents the degree to which the fMRI signal intensities statistically differ between the two levels of the Time variable. For example, in the current picture naming experiment, a high positive t-value at a given voxel and at a given time point indicates that fMRI signal intensities were higher at that time point compared to the baseline values. In our experiment, baseline values corresponded to the fMRI signal intensities observed during time point 1. Thus, there is a difference in interpretation of the extracted BOLD signal by the standard and Slice-Based methods. Whereas in the standard time-series approach the individual t-values indicate the degree of (partial) correlation between the stimulus presentation scheme and the observed BOLD fluctuations across the whole imaging run, the t-values in the time point-by-time point approach directly index differences in measured fMRI signal intensity values between a particular time point in the epoch and the baseline.

The implication of such differences in the statistical modeling methods of fMRI data are currently not well understood. Further research is clearly required. We are currently investigating this issue in terms of the following questions. First, does basic signal detection differ between the time-series and time point-by-time point approaches? What is the effect of using different types of baselines within a time point-by-time point approach (Stark & Squire, 2001)? Can time point-by-time point approaches be applied to blocked designs? Is prewhitening less of an issue in time point-by-time point approaches (e.g., Woolrich et al., 2001)? Can the time point-by-time point approach more easily accommodate new modeling techniques that can handle more complex random effect structures (LME; e.g., Bates, 2005; Pinheiro & Bates, 2000)? Questions such as these should elucidate the exact role of these types of analyses for fMRI research.

### Impact of pre-processing

Current fMRI data analysis packages offer a wide range of different pre-processing options to minimize the impact of noise on statistical modeling. For example, current standard pre-processing tools include, among others, temporal filtering, motion correction, and spatial smoothing. What impact do the different kinds of pre-processing options have on the time course of the BOLD signal extracted by the Slice-Based method? We have investigated this issue to some extent using the data from the picture naming experiment. Note that the data as they were presented above were minimally pre-processed, meaning that only temporal filtering and motion correction tools were applied to the raw data. No smoothing was applied. To examine the impact of various types of pre-processing we created two additional data sets: One in which the raw data was only temporally filtered, and one in which the data was temporally filtered, motion corrected, and 2D smoothed at 5 mm FWHM. We will refer to these three additional data sets as the no, minimally, and maximally pre-processed data sets, respectively. Note that we did not apply any 3D smoothing because this would change signal intensities across slices which would directly affect the temporal accuracy of the signal. All pre-processing tools came from FSL (Jenkinson et al., 2012). We compared these three data sets in terms of the interslice correlation (IC), number of unique peaks (UP), time to peak (TTP), and maximum t-value (MAXT).

We found that the IC differed between the no pre-processing and the maximally pre-processed set (*t*(29) = 2.94, *p* < 0.007), indicating increased interslice correlation in the maximally pre-processed data set compared to the no pre-processing set (0.83 vs 0.89). In addition, the UP differed between the no and the minimally pre-processed set (*t*(29) = 2.41, *p* < 0.03) due to a lower number of peaks in the latter set (1.6 vs 1.5). Similarly, the UP differed between the minimally and the maximally preprocessed set (*t*(29) = 2.25, *p* < 0.04) due to a lower number of peaks in the latter set (1.5 vs 1.4). The TTP differed between the no and the minimally pre-processed set (*t*(29) = 2.28, *p* < 0.03) indicating a later TTP in the minimally pre-processed set (4.1 vs 4.3 s). Furthermore, the maximum t-value of the BOLD time course differed between the not pre-processed and the maximally pre-processed data sets (*t*(29) = 5.13, *p* < 0.001) indicating higher t-values in the maximally pre-processed set (6.5 vs 7.5). Finally, the maximum t-value differed between the minimally and the maximally pre-processed data sets (*t*(29) = 3.97, *p* < 0.001), indicating increased t-values in the latter set (6.6 vs 7.5).

These analyses show that different types of pre-processing tools impacted various measures associated with the time course of the BOLD signal extracted with the Slice-Based method. However, it is important to point out that these analyses do not show the impact of pre-processing on the temporal accuracy of the extracted BOLD signal. In other words, although it is clear that pre-processing tools impacted the time course of the extracted BOLD signal, it remains unclear whether such pre-processing tools actually improve or hamper the temporal accuracy of the BOLD signal extracted with the Slice-Based method. Determining this relationship between different types of pre-processing tools and the Slice-Based method goes beyond the goals of the current study. Future studies that rely on simulated ground-truth signals should be able to resolve this issue.

### Implications

The Slice-Based framework for fMRI data analysis has the ability to extract BOLD signals for the whole-brain with a high temporal accuracy and at a very high temporal resolution. The method has implications for at least three different fields of research. First, in event-related fMRI studies, extracted BOLD signals for different stimulus conditions may be used to make inferences about the underlying neural processes (e.g., Friston et al., 1998; Josephs et al., 1997; Toni et al., 1999). For such studies, extracting a more temporally accurate and higher temporal resolution BOLD signal may enable more fine grained comparisons. Similarly, the improved accuracy and temporal resolution of the Slice-Based method may lead to more accurate functional connectivity maps in task-based functional connectivity studies (Biswal et al., 1995; Rissman et al., 2004). However, both these types of studies should be aware of the limitations of using the BOLD signal to index neural activity. Although many studies now show that BOLD signals index localized neural activity in the brain, the precise dynamics of the BOLD signal has been shown to vary across different regions of the brain, and even within the same region across different participants (e.g., Logothetis & Wandell, 2004). In addition, a substantial body of research has examined the limits to which experimentally induced temporal delays in neural activity can be recovered from the observed fMRI BOLD signal (e.g., Formisano & Goebel, 2003; Katwal et al., 2013; Kim et al., 1997; Kruggel & von Cramon, 1999; Menon et al., 1998; Menon, 2012; Miezin et al., 2000; Ollinger et al., 2001). Thus, although the Slice-Based method can extract BOLD signals at a theoretical maximum temporal resolution that is on the order of tens of milliseconds, researchers should also be aware of the limitations of using the dynamics of the BOLD signal as a proxy for the dynamics of neural activity.

Finally, the current results have implications for studies of the mechanisms underlying neuro-vascular coupling (Attwell & Iadecola, 2002; Hillman, 2014; Logothetis & Wandell, 2004; Uludağ & Uğurbil, 2015). These studies have relied on the BOLD signal to examine the impact of neural activity on vascular changes. These studies have often relied on optical imaging methods to study the temporal dynamics of the BOLD signal (e.g., Chen et al., 2011; Peppiatt et al., 2006). Optical imaging does not have the temporal resolution limitations that are common to MRI. However, optical imaging studies are limited to the study of BOLD signals in the superficial vasculature of the cortex due to the low penetrating depth of light used in the optical techniques. By contrast, the Slice-Based technique permits the simultaneous extraction of highly accurate and high temporal resolution BOLD signals from both cortical and subcortical areas of the brain. This may therefore permit a more general view of of how vascular responses are affected by neural activity.

### Relationship with previous studies

Previous studies in fMRI data analysis have suggested that the temporal resolution of the BOLD signal may be improved by so-called *jittering* of stimulus presentations (e.g., Dale, 1999; Josephs et al., 1997; Price et al., 1999; Toni et al., 1999). In these studies, stimuli are presented at variable time intervals along the course of the experiment. This method of stimulus presentation then allows for FIR basis functions to extract a higher temporal resolution BOLD signal. In the current study, stimulus presentation is also jittered since stimulus presentation must be in-phase with the slice acquisitions. However, the novel aspect of the current study is not that the temporal resolution of the BOLD signal can be improved by a jittered stimulus presentation method. Instead, the novel aspect of the current study lies in the different method of creating whole-brain volumes. A primary requirement of this new method of creating whole-brain volumes is that stimuli must be presented in-phase with the slice-acquisitions (See Figure 2). This volume creation method then leads to whole-brain volumes that contain slices that are all acquired at the same point in time relative to a presented stimulus. A by-product of this method is that the volumes are available at a much higher temporal resolution. Thus, although previous studies have improved the temporal resolution of the BOLD signal by using jittered stimulus presentation, these studies relied on the standard method of volume creation, and therefore, as we have attempted to show in the current study, extracted BOLD signals at a poor temporal accuracy. The novel aspect of the current study is therefore not the improved temporal resolution through the jittered presentation of stimuli, but rather the improved temporal accuracy and temporal resolution of BOLD signal extraction due to the novel method of volume creation.

### Limitations

The slice-based method currently has several limitations. First, the method requires the presentation of a large number of stimuli. Increasing the temporal resolution essentially means distributing a number of collected datapoints across an increasingly larger number of timepoints. This means that in order to attain a sufficient level of statistical power, a sufficiently large number of datapoints are required. This can be achieved by increasing the number of stimuli per participant, or by increasing the number of participants per experiment and doing group-based analyses. The number of available datapoints also immediately bears on the issue of the maximum attainable temporal resolution, because at very large number of timepoints (i.e., a very high temporal resolution), insufficient data may be available for fitting a given statistical model. By lowering the number of timepoints (i.e., a lower temporal resolution), more datapoints per timepoint are available, enabling the fitting of statistical models. Thus, overall, the higher temporal resolution available in the slice-based method requires more stimuli per participant, and more participants per experiment.

Another limitation is that the slice-based framework requires more statistical tests than the standard time-series approach. In the Slice-Based framework, a statistical model is fitted to each voxel in the brain, *at each timepoint*. This means that the number of statistical tests that is performed in the slice-based approach is equal to the number of relevant voxels in the brain times the number of timepoints in the epoch. The large number of statistical tests raises concerns about the issue of multiple comparisons. Although in the current study we did not employ sophisticated correction techniques, we intent to implement more sophisticated corrections in future implementations of the method. One possibility is to expand the current cluster-correction techniques to work with 4D data. Another is to expand methods developed for electrophyiological data to work with fMRI data (Guthrie & Buchwald, 1991; Maris & Oostenveld, 2007).

## Conclusions

To conclude, a well known limitation of the fMRI technique is that fMRI BOLD signals are extracted from the data with poor temporal accuracy and temporal resolution. We have shown that these aspects of fMRI data are in part due to the volume creation method used in current fMRI data analytic approaches. To address these limitations, we have proposed a new Slice-Based data analytic framework that improves the temporal accuracy and temporal resolution of the fMRI BOLD signal. The new framework achieves this improved accuracy and resolution by creating whole-brain volumes that contain slices that are all acquired at the same time point relative to a presented stimulus, and by using non-standard statistical modeling techniques to extract the BOLD signal. We have shown that the new method is more accurate and more precise than the currently available FIR standard methods in the context of both simulated and real-world data. Because this is a new technique, our evaluation was necessarily limited and there are many outstanding questions. However, we think the new technique provides a new alternative to the analysis of fMRI data and may improve our understanding of the neural activity and its associated vascular changes in both health and disease.

## Acknowledgements

This work was supported by The Spanish Ministry of Economy and Competitiveness (RYC2011-08433 and PSI2013-46334 to N.J.). We thank Sara Duque González, Rebeca de Luis Sosa, and Alba Rodríguez González for help with data collection. Correspondence and requests for materials should be addressed to NJ.

